# Aptamer-based DNA nanoswitches for multiplexed protein detection

**DOI:** 10.64898/2026.05.01.722269

**Authors:** Sravya Kovvali, Camryn A. Beckles, Arun Richard Chandrasekaran, Ken Halvorsen

## Abstract

Simple, modular platforms for detecting biologically relevant proteins are critical for applications in clinical diagnostics, healthcare, and research. Here, we have combined aptamer-based protein recognition with our conformationally-responsive DNA nanoswitches to enable simple, sensitive and specific protein detection. We demonstrate dual detection of two clinically relevant blood proteins, thrombin and VEGF as initial proof of concept.

---

Sensitive detection of biologically relevant proteins is crucial for both clinical diagnostics and fundamental research. However, existing assays are typically expensive, time-consuming, or technically complex. Current protein detection methods include classic antibody-based assays such as ELISA^1^ and Western blotting^2^, optical readouts such as fluorescence, chemiluminescence, and surface plasmon resonance^3^, and newer platforms that include electrochemical sensing^4^, nanomaterials^5^, advanced immunosensing^6^, or CRISPR-enabled signal amplification^7^. Many approaches can offer excellent sensitivity and specificity, but often still require specialized instrumentation, surface fabrication, or complex workflows that increase cost and limit accessibility, especially outside of well-equipped laboratories. More recent developments, including multiplexed aptasensors, SERS-based assays, and nanoparticle-assisted platforms, have improved detection limits and turnaround times, but they often remain tied to custom device construction or advanced analytical readouts^8–11^. As a result, there is still strong interest in protein detection methods that combine high sensitivity with simplicity, low cost, and compatibility with standard laboratory infrastructure.

In this work, we demonstrate aptamer-functionalized DNA nanoswitches for protein detection. DNA nanoswitches are self-assembled, conformationally responsive DNA constructs that switch from a linear to looped conformation upon binding a target molecule^12,13^. The conformational change is detectable on an agarose gel where nanoswitches in the linear “off” state migrate faster than those in the looped “on” state. We have used DNA nanoswitches extensively in the past to detect nucleic acid target molecules including oligonucleotides, microRNA, viral RNA, ribosomal RNA, and RNA modifications^14–19^. While the programmable nature of DNA nanoswitches make them amenable to protein detection as well, challenges in antibody-oligonucleotide coupling have limited work in this area. Initially termed NLISA (Nanoswitch-linked immunosorbent assay), researchers have shown DNA nanoswitches to enable sensitive and specific detection of Prostate Specific Antigen (PSA)^20,21^. This method requires protein targets to have a pair of strongly binding antibodies conjugated to oligonucleotides and purified before use. We considered that aptamers could provide a viable alternative to the previously developed workflow. Aptamers are single-stranded DNA or RNA oligos with unique three-dimensional structures that bind with high-affinity to a variety of targets, such as proteins, small molecules, and polymers^22–24^. Aptamers are more stable and cost-effective than antibodies and are also easier to modify, optimize, and integrate into the DNA nanoswitches. In this study, we present our work on incorporating aptamers into DNA nanoswitches to facilitate protein detection.

As a starting point for this work, we chose thrombin as a model protein due to its clinical relevance and the availability of well-validated DNA aptamers. Thrombin is an essential protease in the coagulation cascade (typically in nM to mM concentrations), and its dysregulation is implicated in angiogenesis aberrations^*25*^ and tumor proliferation^*26,27*^. Several thrombin aptamers have been discovered and studied, and we selected two well-established thrombin aptamers from the literature^*28–31*^ (**Table S1**). Thrombin binding aptamer 1, which we designate as TBA1 (**Table S2**), is a 15-nt DNA aptamer that has been shown by NMR to adopt a G-quadruplex structure^*31*^. Thrombin binding aptamer 2 (TBA2) (a 29-nt DNA aptamer) also forms a G-quadruplex structure and binds to the heparin-binding exosite of thrombin^*29*^. These two aptamers have been previously used independently or together for various aptamer-based thrombin sensing schemes^*32–39*^.

To design DNA nanoswitches with thrombin aptamers, we replaced two oligonucleotides spaced ∼2400 nucleotides apart in the structure to include the aptamer sequences (**Figure 1a)**. Oligonucleotides were designed and purchased (IDT DNA) to contain a 40 nt M13 binding region, a 5-thymine spacer for flexibility, and either TBA1 (appended in the 3’ direction) or TBA2 (appended in the 5’ direction) **(Figure 1b)**. When properly integrated into the DNA nanoswitches, these oligos would allow capture and loop formation to detect thrombin.

**Figure 1.**
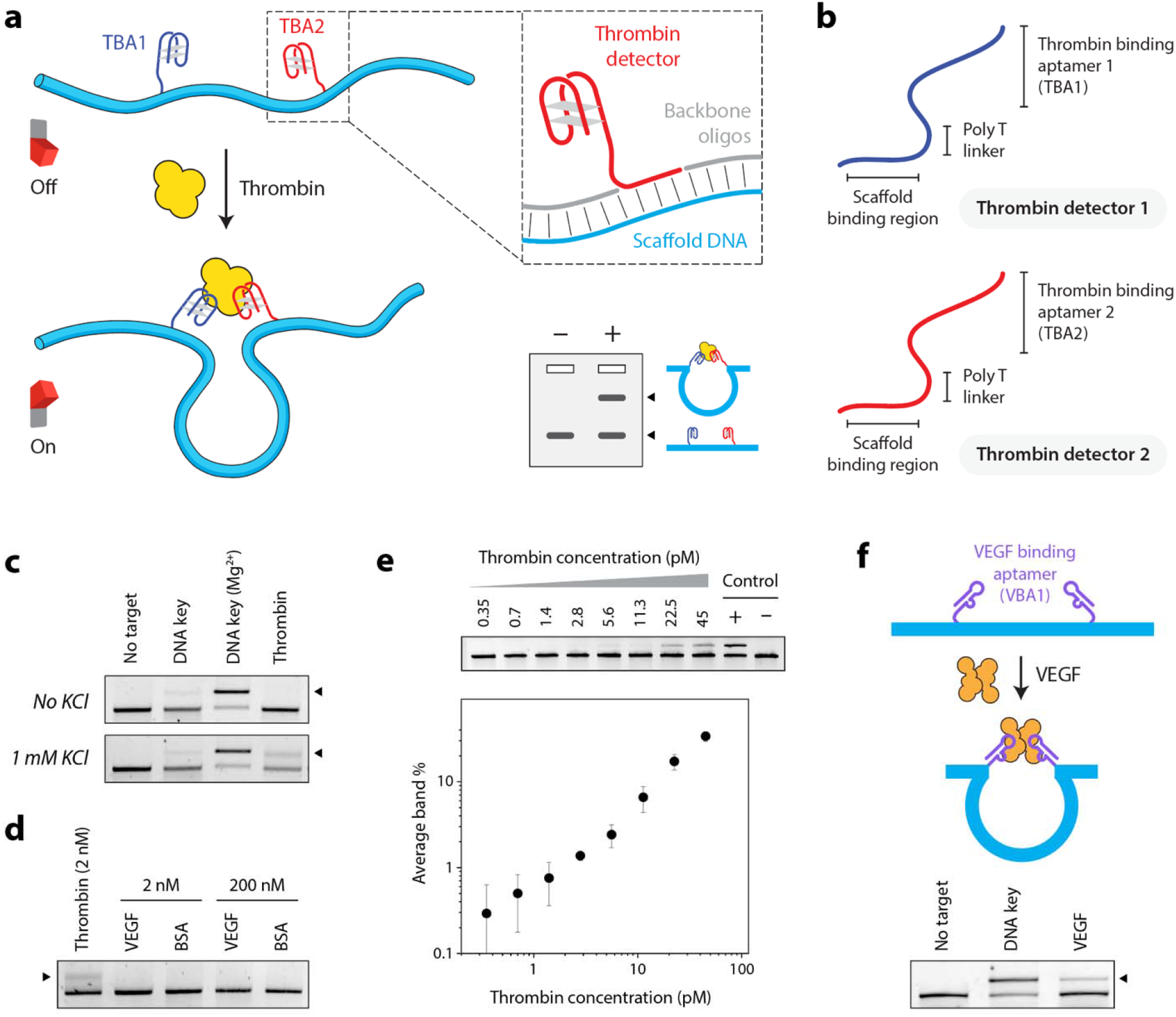
Aptamer-based DNA nanoswitch design and function. (a) Schematic of the thrombin nanoswitch design and operating principle, showing two different thrombin binding aptamers incorporated into the M13 scaffold. (b) Conceptual design of the thrombin detection strands (c) The binding of the aptamers to thrombin requires K+ ions in the incubation buffer as well as the gel and running buffers. (d) Specificity test of thrombin nanoswitch tested against non-target proteins VEGF and BSA at various concentration (e) Sensitivity analysis of thrombin detection showing increased nanoswitch looping with increasing concentrations visualized by agarose gel electrophoresis. (f) Concept and operation of a VEGF binding nanoswitch using aptamers.

As a preliminary test, we sought to confirm thrombin-aptamer binding with these aptamer oligonucleotides, since they necessarily extended the aptamer from its native form to allow nanoswitch integration. We incubated thrombin with each of the thrombin detectors in equimolar concentrations in the “selection buffer” used in their discovery^28^. On a standard TBE agarose gel, we confirmed binding with TBA2 but not TBA1 (**Figure S1a**), leading us to suspect that aptamers may not remain well folded during electrophoresis. When the experiment was repeated with the gel and running buffer spiked with the selection buffer (**Figure S1b**), we confirmed binding of both aptamers. This result suggests that for our nanoswitch assay, both the incubation buffer and the gel running buffer are going to need to be compatible with aptamer folding conditions.

Having established binding with the thrombin detector oligos, we proceeded to test the aptamer-integrated DNA nanoswitches. A short DNA oligonucleotide partly complimentary to each of the aptamers was used as a positive control to confirm the incorporation of both the thrombin detectors into the nanoswitch by hybridizing with each aptamer sequence. We confirmed detection of thrombin using the DNA nanoswitches and established KCl as the critical component of the selection buffer to maintain aptamer binding in the gel (**Figure 1c & S2**). We also demonstrated the specificity of the nanoswitch by evaluating it against 100-fold higher concentrations of off-target proteins (**Figure 1d**), which failed to induce off-target looping of the nanoswitch. Next, we evaluated the sensitivity of the detection by testing various concentrations of thrombin against the nanoswitch, confirming that the nanoswitch could provide visual detection for thrombin concentrations as low as 6 pM (**Figure 1e**). As a whole, these results demonstrate clear proof of concept to integrate aptamers with DNA nanoswitches for sensitive and specific protein sensing with a visual readout.

Following the successful demonstration of nanoswitch-based thrombin detection, we wanted to demonstrate the broad applicability of our approach by detecting a second protein. We chose vascular endothelial growth factor (VEGF) as our second model protein. VEGF is a key regulator of angiogenesis, whose dysregulation is implicated in cardiovascular disease, tumor angiogenesis, and eye diseases including macular degeneration and diabetic retinopathy^40^. While selecting from VEGF aptamers validated in existing literature, it was critical to choose a pair of aptamers that can bind simultaneously to VEGF with sufficient affinity to maintain nanoswitch looping, as some of the validated aptamers did not meet this requirement^41^. A newly developed VEGF aptamer has been shown to bind to VEGF at a ratio of 2:1 (aptamer:VEGF) with a low dissociation constant (K_D_) in the nanomolar range^42^. We chose this aptamer (which we term VBA1) for incorporation into both detectors in the VEGF nanoswitch, following a similar strategy as with thrombin (**Figure 1f and S3**). At this point we also chose to evaluate the effect of PAGE purification of the VEGF detectors. We showed that nanoswitch-based detection of VEGF with the PAGE-purified oligonucleotides improved looping efficiency compared to desalted oligonucleotides (**Figure S3**). This may be due to the sensitivity of aptamer structures to their sequences and decreased binding affinity with truncated sequences that can arise during synthesis.

After successfully confirming the functionality of the chosen VBA1 aptamer, we also assessed other VEGF aptamers previously reported, which we denote VBA2^43^, VBA3^44^ and VBA4^45 39,41–43^. To evaluate the ability of these aptamers to bind to VEGF within the context of the nanoswitch, we constructed nanoswitches with one detector containing the already validated VBA1 aptamer, and the other detector containing one of the four different aptamers (VBA1-4). Unsurprisingly, the validated nanoswitch containing VBA1/VBA1 combination demonstrated the best looping, with only the VBA1/VBA2 showing substantially reduced looping and others not showing detection at all. These results suggest that the VBA1 aptamer has the highest affinity for VEGF, followed by VBA2, then the other two (**Figure S3**).

One of the key features of the DNA nanoswitches is their programmability, which can be used to incorporate various sensing elements at prescribed positions. To further demonstrate this capability in the context of protein detection, we set out to perform dual detection of both thrombin and VEGF. To accomplish dual detection from one sample, we modified the thrombin nanoswitch to create a smaller loop compared to the VEGF nanoswitch by positioning the detectors closer together on the M13 scaffold (**Figure 2a**). By using an equimolar mixture of the VEGF and thrombin nanoswitches, we successfully detect both target proteins either alone or in combination, confirming that each nanoswitch is specific to its respective target and can operate independently (**Figure 2b-c**). These results confirm that our detection approach can be used for detecting multiple different proteins in a single sample using a one-pot reaction.

**Figure 2.**
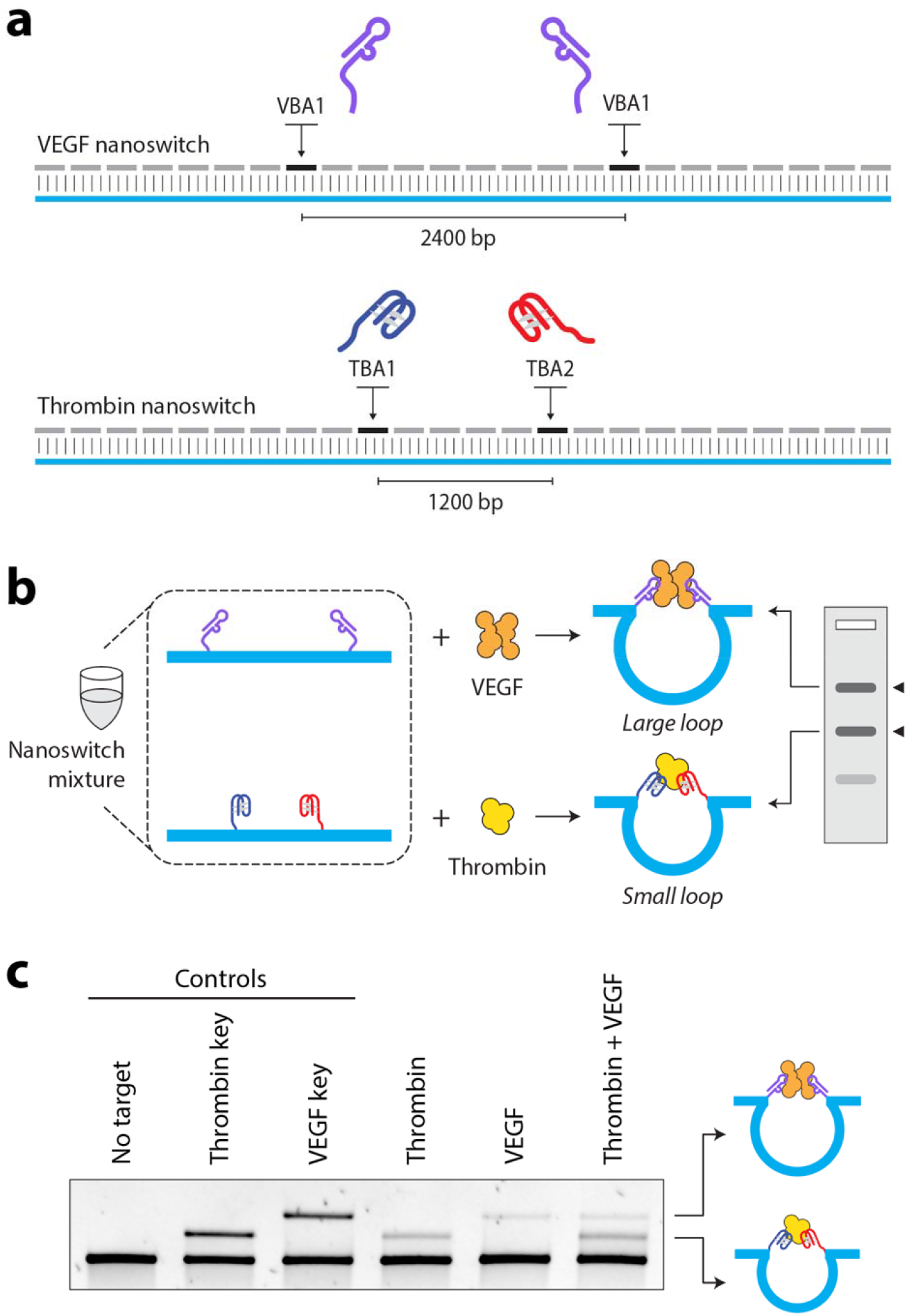
Simultaneous detection of thrombin and VEGF. (a) Schematic of the VEGF and thrombin nanoswitches with different loop sizes. (b) Cartoon of multiplexed composition, workflow, and readout. (c) Agarose gel electrophoresis showing simultaneous detection of thrombin and VEGF from a single sample.

The aptamer-based nanoswitch described in this study provides a simple, modular, and multiplexable platform for protein sensing using the ubiquitous lab method of gel electrophoresis. Our method is perhaps most notable for what it eliminates, enabling specific protein detection entirely without the use of proteins. While most methods rely on antibodies for sensing and either antibodies or enzymes for generating signal, aptamer sensing with DNA nanoswitches eliminates both. Also, unlike many other protein sensing techniques, our method proceeds in a one-step reaction and doesn’t require washing or blotting. The method features visual confirmation of the protein detection, which has long been important for biologists and is likely the primary reason why Western blotting continues to be popular even in the face of newer (arguably better) technologies.

The method does have some caveats and limitations, mainly derived from aptamers. Most obviously, the protein has to maintain binding to two aptamers. Since aptamer-target binding can have a variety of mechanisms (including electrostatic, hydrophobic, etc.) some optimization of buffer and temperature conditions may be required. Furthermore, we have shown that PAGE purification improved aptamer performance (**Figure S4**), highlighting the importance of material quality on aptamer-based detection. Perhaps the most important consideration is aptamer affinity. While it is a good starting point to choose aptamers with low nM reported affinities, we have shown here that published affinity values do not always translate directly to nanoswitch performance. For example, the VEGF aptamers VBA1 and VBA2 have comparable reported K_d_ values (4.0 ± 0.5 nM for VBA1^42^ and 9.9 ± 1.3 nM for VBA2^41^), but only VBA1 produced robust detection. This may suggest that nanoswitch incorporation may alter the aptamer performance, or that K_D_ may not be the correct measure for predicting nanoswitch performance (we suspect the off rate k_off_ may be more important). These factors suggest that aptamer performance may be context dependent and some optimization within the final assay format may be required for new protein targets.

Beyond proteins, our aptamer-based DNA nanoswitch strategy could be expanded for other targets. Several aptamers have been developed for small molecules (e.g. drugs^46^, toxins^47^, etc.), including some that pair a ligand-binding aptamer with a fluorogenic aptamer that activates the fluorescence of a small-molecule dye^48,49^. In principle our strategy may be adopted to some of these cases with further development. Other combined sensing approaches could be used as well, combining aptamers with other sensing modalities to allow multiplexed RNA and protein detection in a single assay. The ability to detect multiple targets using a single electrophoresis readout makes this approach appealing for future diagnostic development, also bolstered by recent developments to streamline the gel process^50^. With further optimization for biological samples and expansion to additional targets, aptamer-based nanoswitches may provide a low-cost and adaptable route to multiplexed biomarker detection.

## Acknowledgements

The authors acknowledge Iranna Todkari, Vinod Morya, Jibin Abraham Punnoose for useful discussions, and Kristina Alejos for early experiments in this area. Research reported in this publication was supported by the National Institutes of Health (NIH) through National Institute of General Medical Sciences (NIGMS) under award number R35GM124720 (to K.H.) and R35GM150672 (to A.R.C.) This manuscript is the result of funding in whole or in part by the National Institutes of Health (NIH). It is subject to the NIH Public Access Policy. Through acceptance of this federal funding, NIH has been given a right to make this manuscript publicly available in PubMed Central upon the Official Date of Publication, as defined by NIH.

## Notes

### Competing Interest Statement

K.H. and A.R.C. have intellectual property related to DNA
nanoswitches. All other authors declare that they have no
competing interests.

