## Supplemental Information for "Aptamer-based DNA nanoswitches for multiplexed protein detection"

**Materials and Methods**

**Linearization of circular M13 DNA**

Single-stranded circular M13 DNA (NEB, Ipswich, MA, USA) was linearized using the BtsCI restriction enzyme (20,000 units/mL, NEB) and a short oligo complementary to the restriction site to facilitate the enzyme activity. The linearization was performed in 1X CutSmart buffer (NEB, Ipswich, MA, USA) at 50°C for 15 minutes in a T100 Thermal Cycler (Bio-Rad, Hercules, CA, USA). The mixture was then heated to 80°C for 20 minutes to deactivate the enzyme. The linearized M13 was stored at 4°C until use.

**Construction of the DNA nanoswitch and purification**

Backbone oligos and variable region tiling oligos (IDT) complementary to the M13 scaffold were added in 10-fold excess to the linearized M13 scaffold. Detector oligos with aptamer sequences (IDT) were purchased with standard desalting or PAGE purification (as noted in the main text) and were also added at roughly 10-fold excess. To accomplish this, all oligos were mixed in equal volumes from 100 µM stocks to make an oligo mixture. The oligo mixture and the linearized M13 were combined in a volume ratio of 1.2:5 for the final construction. The mixture was annealed by ramping down the temperature from 80°C to 12°C at -1°C/min in a T100 Thermal Cycler (Bio-Rad, Hercules, CA, USA). The nanoswitches were purified using the previously established LC purification method. (1)

**Target detection**

Target detection reactions with DNA nanoswitches were carried out in 10 µL volumes. Reactions contained either 1 µL (Figure 1F) or 2 µL (other figures) LC-purified nanoswitch, either 2 µL of VEGF buffer or 1 µL of thrombin buffer, and 1 µL of target protein (or water for negative control), and water to make up the remaining volume to 10 µL. For the VEGF nanoswitches, the VEGF buffer was 5X PBS (Invitrogen, Thermo Fisher Scientific, Carlsbad, CA, USA) spiked with 100 mM KCl and 1 mg/mL BSA. For the thrombin nanoswitches, the thrombin buffer contained 40 mM NaCl2, 20 mM TAE, 5 mM KCl, 1 mM MgCl2, 1 mM CaCl2, and 0.1 mg/mL BSA. Samples were incubated for 1 hour at 37°C for VEGF detection or RT for thrombin detection.

**Gel electrophoresis**

Following incubation, 2 µL of 6X loading dye was mixed with the samples before loading onto 0.8% or 1% agarose (Sigma) gels. The agarose gels were prepared with 0.5X TBE (made in-house) and were supplemented with 1 mM KCl (for thrombin NS gels) or 10 mM KCl (for VEGF nanoswitch gels) unless indicated otherwise. Gels were run for 30 minutes at 75V (for VEGF) or at 25V for 3 hours at 4°C (for thrombin) and post-stained with 1X GelRed (Biotium, Fremont, CA, USA) solution in 100 mM NaCl for 20-30 minutes. The stained gels were imaged using Azure Biosystems (Dublin, CA, USA) imager and analyzed using ImageJ software.

Table S1: Oligonucleotides used in the study. The aptamer portion is indicated in bold.

| Name | Sequence (5’ 🡪 3’) |
| --- | --- |
| Thrombin detector 1 | CAATACTTCTTTGATTAGTAATAACATCACTTTTT**GGTTGGTGTGGTTGG** |
| Thrombin detector 2 | **AGTCCGTGGTAGGGCAGGTTGGGGTGACT**TTTTTTCAACCGATTGAGGGAGGGAAGG |
| Thrombin DNA key | AGTCACCCCAACCTGCCCTACCACGGACTCCAACCACACCAACC |
| VEGF detector 1 | CAATACTTCTTTGATTAGTAATAACATCACTTTTT**CTAGGGGTCCAGGCGAAGCTTAGTAGGGGTGTCCCCTCCCA** |
| VEGF detector 2 | **CTAGGGGTCCAGGCGAAGCTTAGTAGGGGTGTCCCCTCCCA**TTTTTTCAACCGATTGAGGGAGGGAAGG |
| VEGF detector 3 | CAATACTTCTTTGATTAGTAATAACATCACTTTTT**GCCCGTCTTCCAGACAAGAGTGCAGGGC** |
| VEGF detector 4 | CAATACTTCTTTGATTAGTAATAACATCACTTTTT**CAATTGGGCCCGTCCGTATGGTGGGT** |
| VEGF detector 5 | CAATACTTCTTTGATTAGTAATAACATCACTTTTT**TGTGGGGGTGGACTGGGTGGGTACC** |
| VEGF DNA key VBA1-1 | CCCCTACTAAGCTTCGCCTGGACCCCTAGTGGGAGGGGACACCCCTACTAAGCTTCGCC |
| VEGF DNA key VBA 2-1 | CACCCCTACTAAGCTTCGCCTGGACCCCTAGGCCCTGCACTCTTGTCTGGAAGACGGGC |
| VEGF DNA key VBA 3-1 | GACACCCCTACTAAGCTTCGCCTGGACCCCTAGACCCACCATACGGACGGGCCCAATTG |
| VEGF DNA key VBA 4-1 | GGACACCCCTACTAAGCTTCGCCTGGACCCCTAGGGTACCCACCCAGTCCACCCCCACA |

Table S2: Aptamer nomenclature used in the study and corresponding alternatives reported in the literature

| Aptamer name used in this study | Nomenclature reported in literature | Reference |
| --- | --- | --- |
| TBA1 | TBA15, TBA, 15-mer TBA | 28 |
| TBA2 | HD22T5, 29-mer TBA | 29 |
| VBA1 | H4 | 42 |
| VBA2 | 5’-GC-, 3’-C, A | 43 |
| VBA3 | SL2-B, C | 44 |
| VBA4 | 3R02, B | 45 |

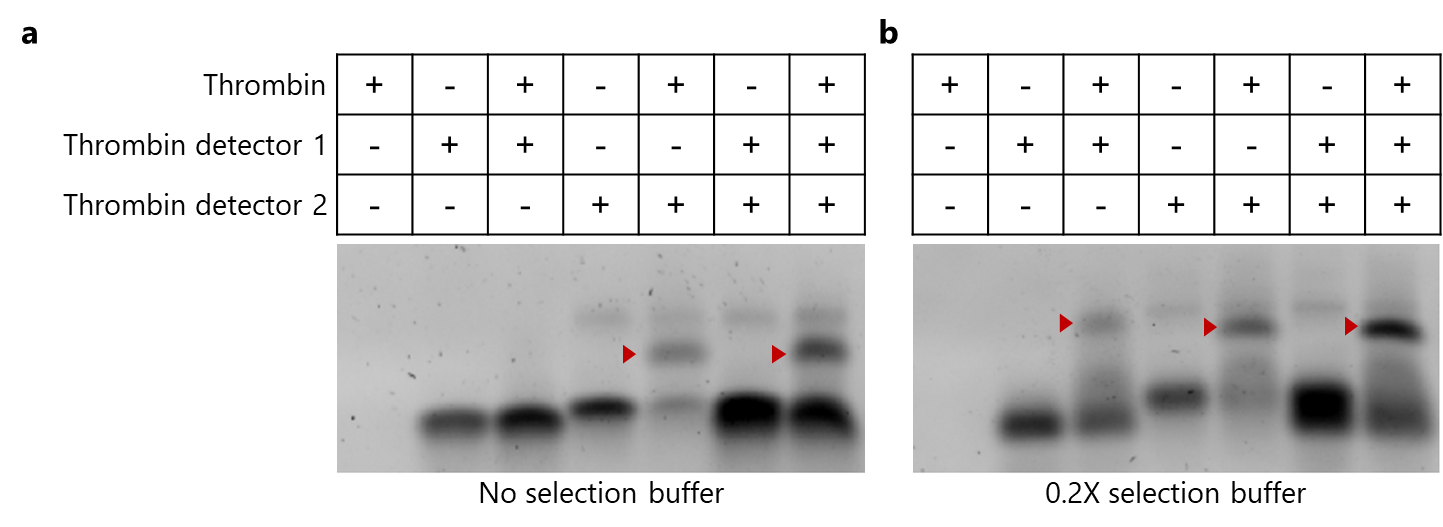

Figure S1: Agarose gel analysis to assess the binding of thrombin to its aptamers. Thrombin and aptamers were at a final concentration of 1 µM. (a) Samples run on a standard 0.5X TBE gel (b) Samples run on a 0.5X TBE gel spiked with selection buffer (0.2X final strength).

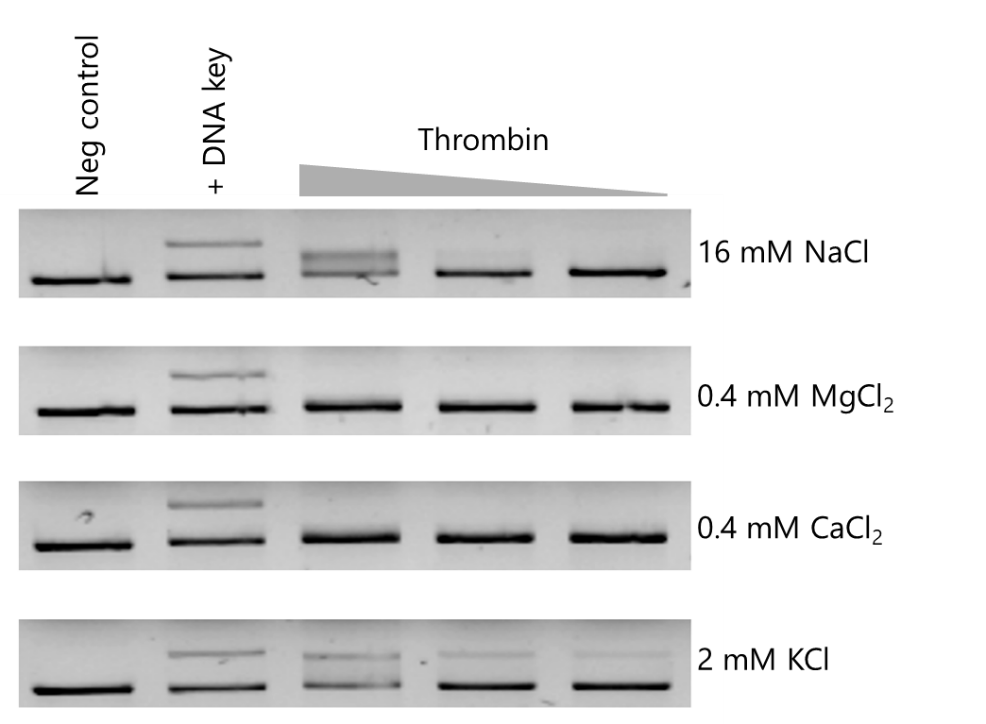
Figure S2: Agarose gel analysis of thrombin nanoswitch at varying thrombin concentrations. The 0.5X TBE gel was supplemented with the indicated selection-buffer component to assess its essentiality for the aptamers binding to thrombin. Nanoswitch looping indicates binding of both the aptamers to the same thrombin molecule.

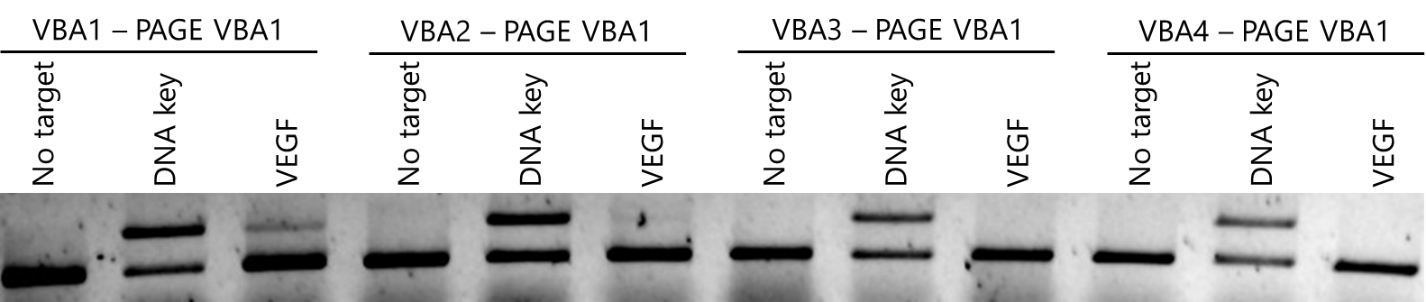

Figure S3: Comparison of nanoswitch performance using different VEGF aptamer pairs. Nanoswitches with one detector containing the PAGE-purified VBA1 aptamer, and the other detector containing one of the four different non-PAGE purified aptamers (VBA1-4).

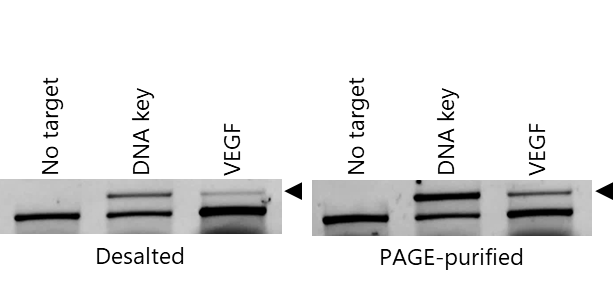

Figure S4: Comparison of looping efficiency of the nanoswitch constructed with desalted and PAGE-purified VEGF detectors.
